# Modeling Retinal Ganglion Cell Degeneration from Longitudinal Intraocular Pressure Trajectories

**DOI:** 10.64898/2026.08.09.743809

**Authors:** Andrew Lesniewski, Margaret A. MacNeil

## Abstract

Many experimental studies collect longitudinal physiological measurements while assessing irreversible biological outcomes only at a terminal endpoint, leaving the timing of disease progression unobserved. This disconnect between continuously measured covariates and latent biological events limits quantitative analysis of how physiological stress drives tissue degeneration. We address this problem by formulating retinal ganglion cell (RGC) degeneration in experimental glaucoma as a latent time-to-event process driven by longitudinal intraocular pressure (IOP) exposure.

Using monthly IOP measurements and terminal RGC counts from the DBA/2J mouse model of glaucoma, we develop both Cox proportional hazards models and a time-dependent extension based on the Andersen–Gill counting-process formulation, allowing progression risk to depend on both contemporaneous IOP and cumulative pressure burden. We further reconstruct model-implied survival curves from the fitted hazard functions, providing a continuous-time representation of latent disease progression under observed and hypothetical IOP trajectories.

Across all disease thresholds and both modeling approaches, cumulative IOP burden above 19 mmHg emerged as the dominant predictor of RGC degeneration, whereas peak and contemporaneous IOP contributed little additional predictive information once sustained exposure was taken into account. HDAP2, a mitochondria-targeted neuroprotective peptide, significantly reduced progression hazard after adjustment for longitudinal IOP exposure, supporting a pressure-independent neuroprotective mechanism.

Beyond identifying cumulative pressure exposure as the dominant predictor of neurodegeneration in this experimental model, the proposed framework provides a general strategy for relating longitudinal physiological measurements to latent biological progression. By linking exposure histories to model-implied survival trajectories, it enables trajectory-based risk assessment, prediction under hypothetical IOP trajectories, and quantitative evaluation of therapeutic interventions in experimental systems where biological outcomes are observed only at terminal endpoints.

## 1 Introduction

Survival analysis is traditionally formulated for settings in which the time of a biological event is observed directly [1]. In many experimental systems, however, the opposite situation arises: physiological variables can be measured repeatedly throughout the course of disease, whereas the biological event of interest is observed only indirectly through a terminal endpoint. This disconnect between longitudinal covariates and latent event times limits our ability to reconstruct disease progression and to determine how evolving physiological stress influences irreversible tissue degeneration. Developing statistical frameworks that bridge this gap remains an important challenge in quantitative biology.

Experimental glaucoma provides a compelling example of this problem. In the DBA/2J mouse model, intraocular pressure (IOP) can be measured longitudinally over many months, providing a detailed longitudinal record of mechanical stress experienced by each eye [26]. In contrast, retinal ganglion cell (RGC) survival is determined only at sacrifice by counting immunolabeled cells in retinal wholemounts [17]. The pressure history of every animal is therefore known, but the time at which biologically meaningful neurodegeneration occurred remains unobserved. Two animals sacrificed at the same age may exhibit similar terminal RGC counts despite having experienced markedly different patterns of IOP elevation, while animals with comparable cumulative pressure exposure may differ substantially in the timing of disease onset. Extracting dynamic information from this type of dataset requires a framework that explicitly links longitudinal exposure histories to latent disease progression.

Most previous quantitative analyses of glaucoma progression have summarized longitudinal IOP trajectories using a small number of scalar descriptors, including mean IOP, peak IOP, pressure fluctuation, or cumulative pressure burden, and related these measures to structural or functional outcomes [13], [18], [20], [7], [24]. These approaches have established elevated IOP as the principal modifiable risk factor for glaucoma progression and have identified several features of the pressure history that correlate with disease severity [4], [20]. However, reducing an entire trajectory to a few summary statistics necessarily discards information about when pressure elevations occurred and how they accumulated over time. Likewise, previous applications of survival analysis in glaucoma have generally relied on baseline or time-averaged covariates, while machine-learning approaches have emphasized prediction rather than mechanistic interpretation [10], [12], [13], [24], [25]. Joint longitudinal-survival models provide an alternative strategy but are primarily designed for settings with noisy, irregularly sampled biomarkers rather than the controlled longitudinal measurements characteristic of experimental glaucoma [23].

In this study, we formulate RGC degeneration as a latent time-to-event process driven by an animal’s longitudinal IOP history. Using longitudinal IOP measurements and terminal RGC counts from the DBA/2J mouse model [17], we develop both conventional Cox proportional hazards models and a time-dependent extension based on the Andersen-Gill counting-process formulation [3], [1], allowing progression risk to depend on both contemporaneous IOP and cumulative pressure burden. We then reconstruct continuous-time, model-implied survival curves from the fitted hazard functions, providing an explicit representation of latent disease progression and enabling prediction under observed as well as hypothetical IOP trajectories.

We apply the framework to a longitudinal dataset generated during evaluation of HDAP2, a mitochondria-targeted neuroprotective peptide that preserves retinal ganglion cells without altering intraocular pressure [17]. This provides an opportunity not only to identify the features of pressure history that best predict neurodegeneration but also to determine whether neuroprotection remains significant after accounting for the complete longitudinal history of IOP exposure. Because the model explicitly separates pressure-dependent and pressure-independent effects, it also provides a quantitative framework for evaluating neuroprotective therapies that act downstream of ocular hypertension.

The principal contributions of this study are fourfold. First, we develop a survival-analysis framework for experimental systems in which longitudinal physiological measurements are available but event times remain latent. Second, we demonstrate that cumulative IOP burden, rather than peak or contemporaneous IOP, is the dominant predictor of retinal ganglion cell degeneration across multiple disease thresholds. Third, we show that HDAP2 significantly reduces progression hazard even after adjustment for longitudinal pressure exposure, supporting a pressure-independent neuroprotective mechanism. Finally, we introduce a practical method for reconstructing model-implied survival trajectories from longitudinal physiological data, enabling prediction under hypothetical IOP trajectories, and quantitative evaluation of therapeutic interventions.

The remainder of the paper is organized as follows. Section 2 describes the experimental background and data provenance. Section 3 describes the structure of the dataset and the definition of RGC-based endpoints. Section 4 develops the baseline proportional hazards model using static covariates. Section 5 introduces the time-dependent Cox model and its counting-process formulation. Section 6 presents the methodology for constructing model-implied survival curves.

## 2 Animal Methods

The analyses presented here are based on a previously published longitudinal study of retinal ganglion cell (RGC) neuroprotection in the DBA/2J mouse model of glaucoma [17]. Understanding the experimental design is essential because it determines the statistical structure of the dataset and motivates the survival-analysis framework developed in the following sections. In particular, each animal contributes a longitudinal history of intraocular pressure (IOP) measurements together with a single terminal assessment of RGC survival, creating a setting in which physiological covariates are observed longitudinally while the biological event of interest is observed only at the experimental endpoint.

### 2.1 The DBA/2J Mouse Model of Glaucoma

DBA/2J mice develop spontaneous ocular hypertension as a consequence of progressive anterior-segment pathology caused by recessive mutations in *Gpnmb* and *Tyrp1* [2]. IOP typically begins to rise between 6 and 9 months of age, although both the timing and magnitude of pressure elevation vary substantially among individuals [15]. As a consequence, animals accumulate markedly different pressure histories despite being maintained under identical experimental conditions. This heterogeneity is a defining feature of the dataset rather than a source of experimental noise. Approximately 26% of DBA/2J eyes never develop sustained ocular hypertension [16] and consequently remain largely protected from pressure-induced RGC degeneration throughout the observation period. These animals contribute low-exposure trajectories that remain right-censored across all degeneration endpoints, providing an important reference against which the effects of increasing cumulative pressure burden can be estimated. The broad spectrum of naturally occurring IOP trajectories therefore provides the variation required to relate longitudinal pressure exposure to subsequent neurodegeneration.

### 2.2 Animal Experiments

Animals were assigned at 4 months of age to either an untreated control group or a group receiving HDAP2, a mitochondria-targeted neuroprotective peptide administered intraperitoneally every other day. IOP was measured monthly under light isoflurane anesthesia from 4 through 12 months of age using rebound tonometry, providing a longitudinal record of IOP exposure for each animal. At sacrifice, RGC survival was quantified by immunolabeling retinal wholemounts for the RGC-specific marker RBPMS and counting labeled cells with the validated deep-learning platform RGCode, as previously described [17].

Animals were euthanized at 10, 11, or 12 months of age according to the original experimental design [17]. Because sacrifice occurred at multiple ages, the resulting dataset spans a broad range of disease severity and cumulative IOP exposure, providing the variation required for the survival analysis developed below.

### 2.3 RGC Counts and Their Distribution

Healthy DBA/2J retinas before the onset of ocular hypertension contain approximately 50,000 RBPMS-positive RGCs [9]. As disease progresses, terminal RGC counts decline, but the distribution of counts is strikingly non-uniform [14]. Most animals either retain relatively high numbers of RGCs (*>* 38, 000 cells) or exhibit advanced degeneration (*<* 18, 000 cells), with comparatively few intermediate cases.

This distribution has important consequences for endpoint definition. Thresholds placed within the sparsely populated intermediate region are statistically unstable because relatively few animals cross them. Accordingly, the principal analyses emphasize thresholds that coincide with the natural structure of the data. The earliest detectable deficit EDD_75_ (earliest detectable deficit, *≈* 25% loss) marks the transition out of the high-count population, whereas aNLP_35_ (advanced degeneration) represents advanced neurodegeneration. Together these endpoints provide the most biologically informative and statistically robust characterization of disease progression in this cohort.

## 3 Data Structure and Endpoint Definition

The experimental design described in Section 2 produces an animal-level dataset in which longitudinal physiological measurements are paired with a single terminal measure of neurodegeneration. This structure is central to the statistical framework developed below: while IOP is observed repeatedly throughout the experiment, RGC degeneration is observed only at sacrifice, so the time at which a degeneration threshold is crossed remains unobserved.

### 3.1 Dataset Construction

The dataset consists of an animal-level table containing:

1. a unique identifier (animal id),
2. age at sacrifice in months (age months),
3. terminal RGC count (rgc count),
4. treatment condition (Condition *∈ {*untreated, HDAP2*}*),
5. longitudinal IOP measurements at months *t ∈ {*4, 5, 6, 7, 8, 9, 10, 11, 12*}*.

Each row corresponds to a single animal and represents its longitudinal IOP trajectory in *wide format*,

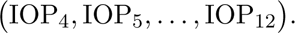

The analysis is restricted to animals sacrificed at or before 12 months of age because HDAP2-treated animals are available only through this time point. Sex is not included as a covariate in the present analysis.

Although IOP is measured longitudinally, RGC counts are obtained only once, at sacrifice. Consequently, the exact timing of biologically meaningful degeneration is not observed directly but must be inferred from the combination of longitudinal pressure history and terminal outcome. This mismatch between covariate and outcome is the principal motivation for the survival modeling framework developed in the following sections.

Missing or non-numeric IOP values are treated as NaN and excluded when constructing derived covariates. All IOP-based quantities are computed using measurements strictly prior to the animal’s sacrifice age.

### 3.2 Event Definitions via RGC Thresholds

RGC degeneration is characterized using four thresholds spanning the range of disease severity observed in this cohort. The thresholds are expressed as fractions of a nominal healthy baseline of 50,000 RGCs, consistent with counts observed in young DBA/2J mice before the onset of ocular hypertension and with previous studies relating RGC loss to functional impairment and disease stage [22], [8], [5]:

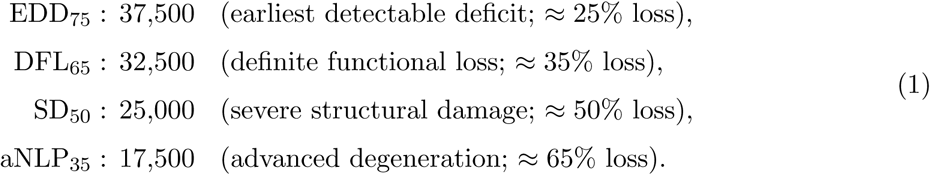

The primary endpoint throughout this study is DFL_65_, which represents a biologically meaningful level of RGC loss associated with measurable functional impairment [11]. Results are presented for all four thresholds to evaluate the robustness of the modeling framework across multiple stages of disease progression.

As discussed in §2.3, the observed distribution of terminal RGC counts is strongly non-uniform. Intermediate thresholds, particularly SD_50_, are crossed by relatively few animals and therefore produce wider confidence intervals than the early and late endpoints. These results are included for completeness but are interpreted with appropriate caution.

For each animal *i* and threshold *c*, define the event indicator

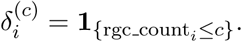

Let *T_i_* = age months denote the observed time. For animals with rgc count*_i_ ≤ c* we assign the event time as *T_i_*, the age at sacrifice; animals with rgc count*_i_ > c* are right-censored at *T_i_* [1]. Because the actual time of threshold crossing is unobserved, *T_i_* represents the terminal observation time at which threshold crossing is known to have occurred, rather than the exact biological event time.

### 3.3 Cross-Sectional Degeneration Frequencies at Sacrifice

Because RGC counts are available only at the terminal sacrifice ages (10, 11, and 12 months), conventional nonparametric survival estimators such as Kaplan–Meier curves [1] cannot be constructed directly. Instead, we first summarize the observed data using cross-sectional degeneration frequencies.

For each degeneration threshold, we compute the proportion of eyes that have reached the corresponding endpoint at each sacrifice age, separately for untreated and HDAP2-treated animals. These summaries provide a direct, assumption-free description of disease burden prior to introducing the survival models.

Several consistent patterns emerge across thresholds. First, HDAP2-treated animals exhibit systematically lower degeneration frequencies than untreated animals at all observed ages. This difference is already apparent at the 10-month time point and becomes more pronounced at later ages, indicating a sustained protective effect.

Second, at 10 months, degeneration is frequently observed in untreated animals but is markedly reduced or absent in the HDAP2-treated group. This suggests that HDAP2 delays the onset of structural degeneration in addition to reducing its overall prevalence.

Third, the separation between treatment groups increases with the severity of the degeneration threshold, consistent with cumulative divergence in disease trajectories over time.

These descriptive summaries motivate the multivariable survival analyses developed in the following sections, which quantify treatment effects while accounting for differences in longitudinal IOP exposure.

## 4 Proportional Hazards Modeling

Having defined the experimental endpoints and the structure of the dataset, we first consider a conventional Cox proportional hazards model in which each animal’s longitudinal IOP history is summarized by a small set of biologically motivated covariates. This provides a baseline for evaluating the relationship between pressure exposure and RGC degeneration and for assessing the efficacy of HDAP2 after adjustment for IOP-derived factors, before introducing the time-dependent framework developed in Section 5.

### 4.1 Static Covariates

For each animal *i*, let

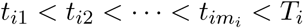

denote the times of its *m_i_* valid IOP measurements strictly prior to sacrifice at age *T_i_*, and let

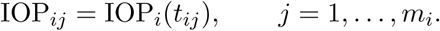

Each animal’s longitudinal IOP trajectory is summarized by the following covariates:

1. *Maximum IOP*

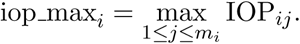

1. *IOP slope*

The longitudinal trend in IOP is represented by the slope obtained from the least-squares fit IOP*_ij_ ≈ a_i_* + *b_i_*(*t_ij_ − t_i_*_1_)*, j* = 1*,…, m_i_,*

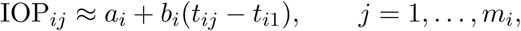

with

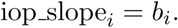

1. *IOP burden above 19 mmHg*

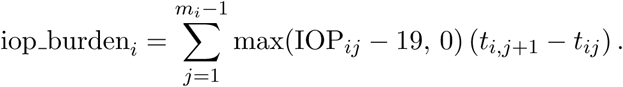

This quantity provides a discrete approximation to cumulative excess pressure exposure above 19 mmHg. The integral is approximated using a piecewise left-constant representation of IOP, consistent with the time-dependent construction in Section 5.

1. *Treatment indicator*

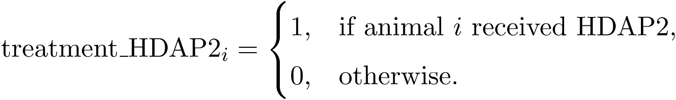

We collect these features into the covariate vector

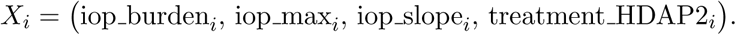

These covariates capture complementary aspects of IOP exposure that are biologically motivated by prior experimental work. Cumulative burden above 19 mmHg measures the integrated physiological stress imposed by sustained pressure elevation rather than any single IOP measurement. Maximum IOP captures the peak pressure experienced by the eye, while IOP slope characterizes the longitudinal trend in pressure over the observation window. The 19 mmHg threshold follows [17], where it was adopted based on established cutoffs in the DBA/2J literature for defining pathologically elevated IOP, and corresponds approximately to the upper bound of normal IOP measured by rebound tonometry in young DBA/2J mice prior to pigment-dispersion onset [16].

### 4.2 Cox Proportional Hazards Model

For each RGC threshold *c*, we model the hazard of progression using the standard proportional-hazards specification [1] as

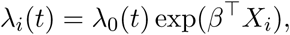

where *λ*_0_(*t*) is a baseline hazard common to all animals, and where *β* is the vector of covariate loadings.

The model is estimated via partial likelihood [1] using the Python lifelines library [6]. Treatment is included explicitly as a covariate, enabling direct inference on HDAP2 efficacy.

### 4.3 Results

Table 1 reports the estimated regression coefficients, standard errors, *p*-values, hazard ratios, and corresponding 95% confidence intervals for each endpoint.

**Table 1:** Cox proportional hazards regression results across RGC loss thresholds. Hazard ratios (HR) are reported as exp(*β̂*), with 95% confidence intervals in parentheses.

| Covariate | $\hat{\beta}$ | SE | $p$-value | HR | 95% CI |
| --- | --- | --- | --- | --- | --- |
| **EDD_75** |  |  |  |  |  |
| iop_burden_19 | 0.120 | 0.045 | 0.00773 | 1.13 | (1.03, 1.23) |
| iop_max | -0.066 | 0.081 | 0.415 | 0.94 | (0.80, 1.10) |
| iop_slope | 0.361 | 0.300 | 0.229 | 1.43 | (0.80, 2.58) |
| treatment_HDAP2 | -1.490 | 0.514 | 0.00377 | 0.23 | (0.08, 0.62) |
| **DFL_65** |  |  |  |  |  |
| iop_burden_19 | 0.140 | 0.047 | 0.00272 | 1.15 | (1.05, 1.26) |
| iop_max | -0.107 | 0.084 | 0.203 | 0.90 | (0.76, 1.06) |
| iop_slope | 0.416 | 0.319 | 0.193 | 1.52 | (0.81, 2.83) |
| treatment_HDAP2 | -1.746 | 0.555 | 0.00164 | 0.17 | (0.06, 0.52) |
| **SD_50** |  |  |  |  |  |
| iop_burden_19 | 0.126 | 0.051 | 0.0143 | 1.13 | (1.03, 1.25) |
| iop_max | -0.066 | 0.091 | 0.472 | 0.94 | (0.78, 1.12) |
| iop_slope | 0.323 | 0.341 | 0.342 | 1.38 | (0.71, 2.69) |
| treatment_HDAP2 | -1.688 | 0.589 | 0.00412 | 0.18 | (0.06, 0.59) |
| **aNLP_35** |  |  |  |  |  |
| iop_burden_19 | 0.146 | 0.053 | 0.00578 | 1.16 | (1.04, 1.28) |
| iop_max | -0.086 | 0.096 | 0.368 | 0.92 | (0.76, 1.11) |
| iop_slope | 0.610 | 0.364 | 0.0934 | 1.84 | (0.90, 3.75) |
| treatment_HDAP2 | -1.634 | 0.606 | 0.00695 | 0.20 | (0.06, 0.64) |

The Cox model leaves the baseline hazard function *λ*_0_(*t*) unspecified and estimates it nonparametrically from the data [1]. The estimated cumulative baseline hazard Λ̂_0_(*t*) provides the anchor for translating relative risk estimates into absolute survival curves, as developed in Section 6.

Cumulative IOP burden above 19 mmHg emerges as the dominant predictor of RGC degeneration across all four endpoints. The estimated coefficients were positive, statistically significant, and stable across thresholds, indicating that integrated pressure exposure provides strongest predictive information. This finding is consistent with longitudinal PERG studies in DBA/2J mice, which showed that RGC functional decline tracks cumulative IOP exposure rather than instantaneous pressure [21], and that the reversibility of pressure-induced dysfunction diminishes with sustained exposure [19]. The present analysis extends these observations by incorporating cumulative burden, peak IOP, pressure trajectory, and treatment within a single Cox model.

In contrast, peak IOP and IOP slope exhibited weaker and less consistent effects across thresholds. Once sustained exposure was included in the model, peak IOP did not provide significant additional predictive information.

The treatment indicator showed a strong and statistically significant protective effect across all endpoints, with HDAP2-treated animals exhibiting substantially reduced hazard of reaching each RGC loss threshold. Importantly, this effect persisted after adjustment for cumulative IOP burden. This finding is consistent with the previously reported mechanism of HDAP2, which stabilizes the inner mitochondrial membrane and protects RGCs from the downstream consequences of sustained IOP stress [17].

### 4.4 Model-Implied Survival Curves from the Proportional Hazards Model

To translate the proportional hazards model estimates into a more interpretable form, we construct model-implied survival curves under representative covariate configurations. These curves combine the estimated regression coefficients with the nonparametric estimate of the baseline hazard to yield predicted survival probabilities as a function of time.

Figure 5 illustrates the model-implied survival curves across all RGC loss thresholds, comparing untreated and HDAP2-treated animals under a fixed set of covariates. Cumulative IOP burden, peak IOP, and slope are held constant, while treatment assignment is varied.

**Figure 1:**
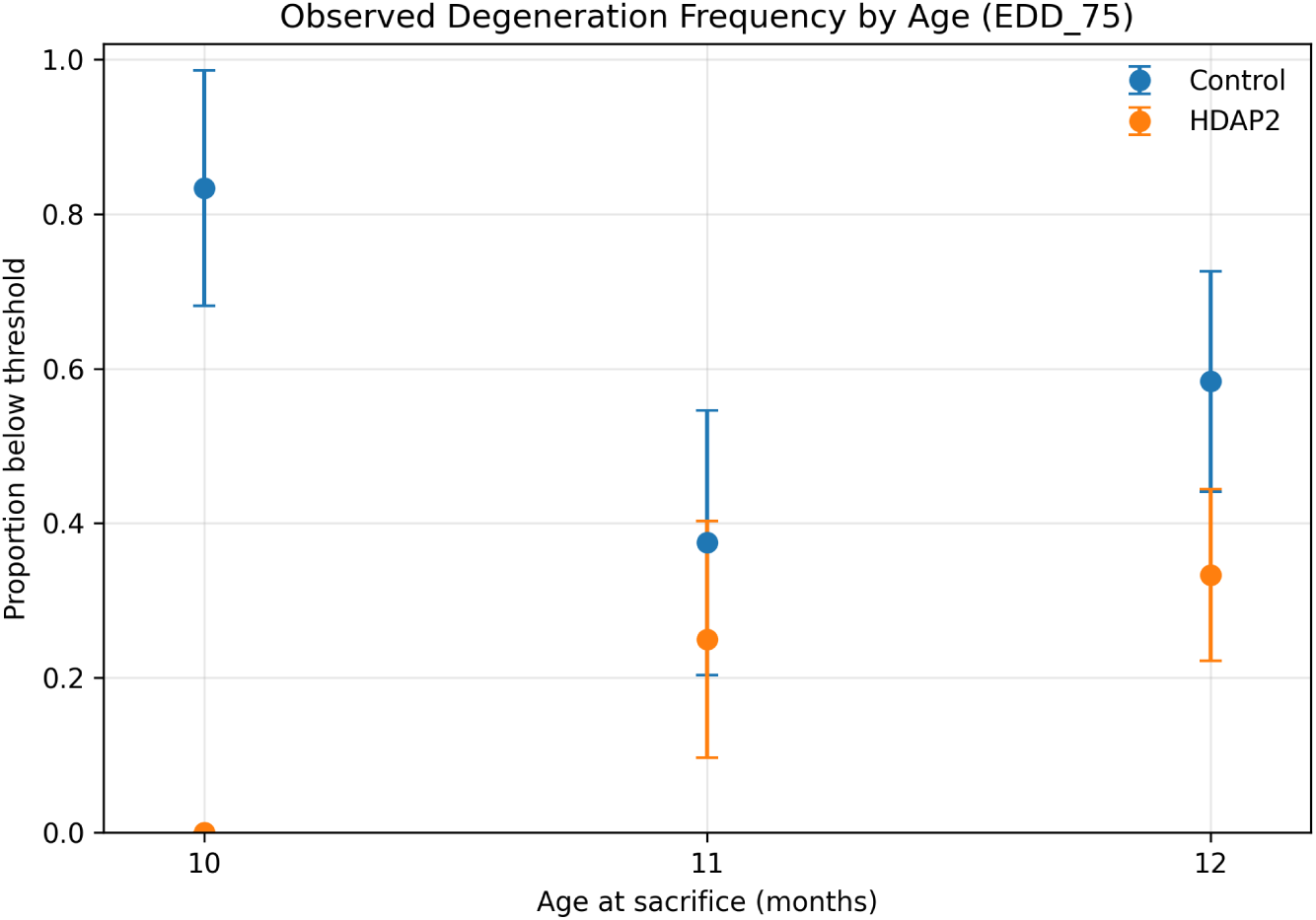
Observed degeneration frequency at the EDD_75_ threshold (earliest detectable deficit) as a function of age at sacrifice. Error bars represent binomial standard errors.

**Figure 2:**
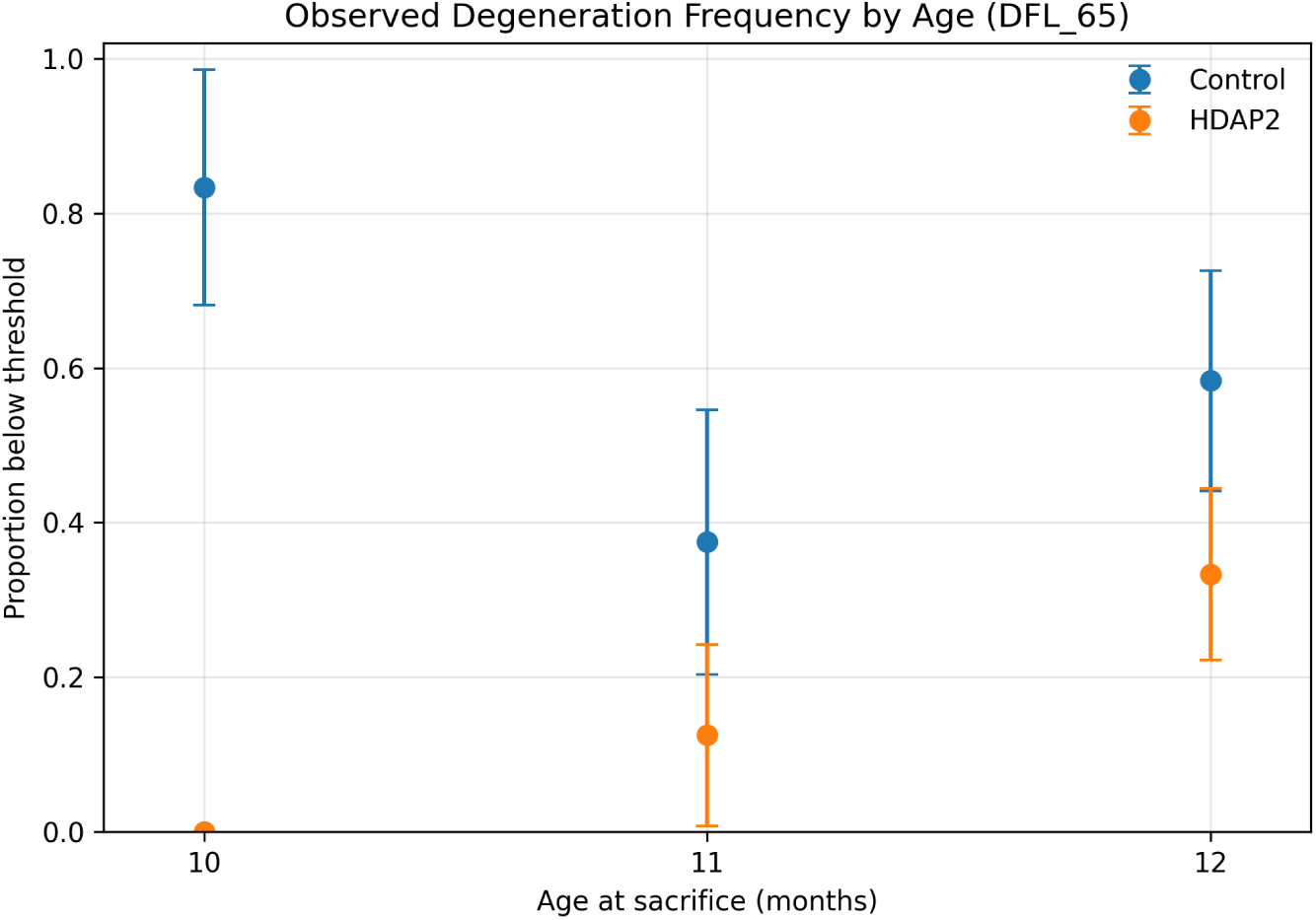
Observed degeneration frequency at the DFL_65_ threshold (definite functional loss).

**Figure 3:**
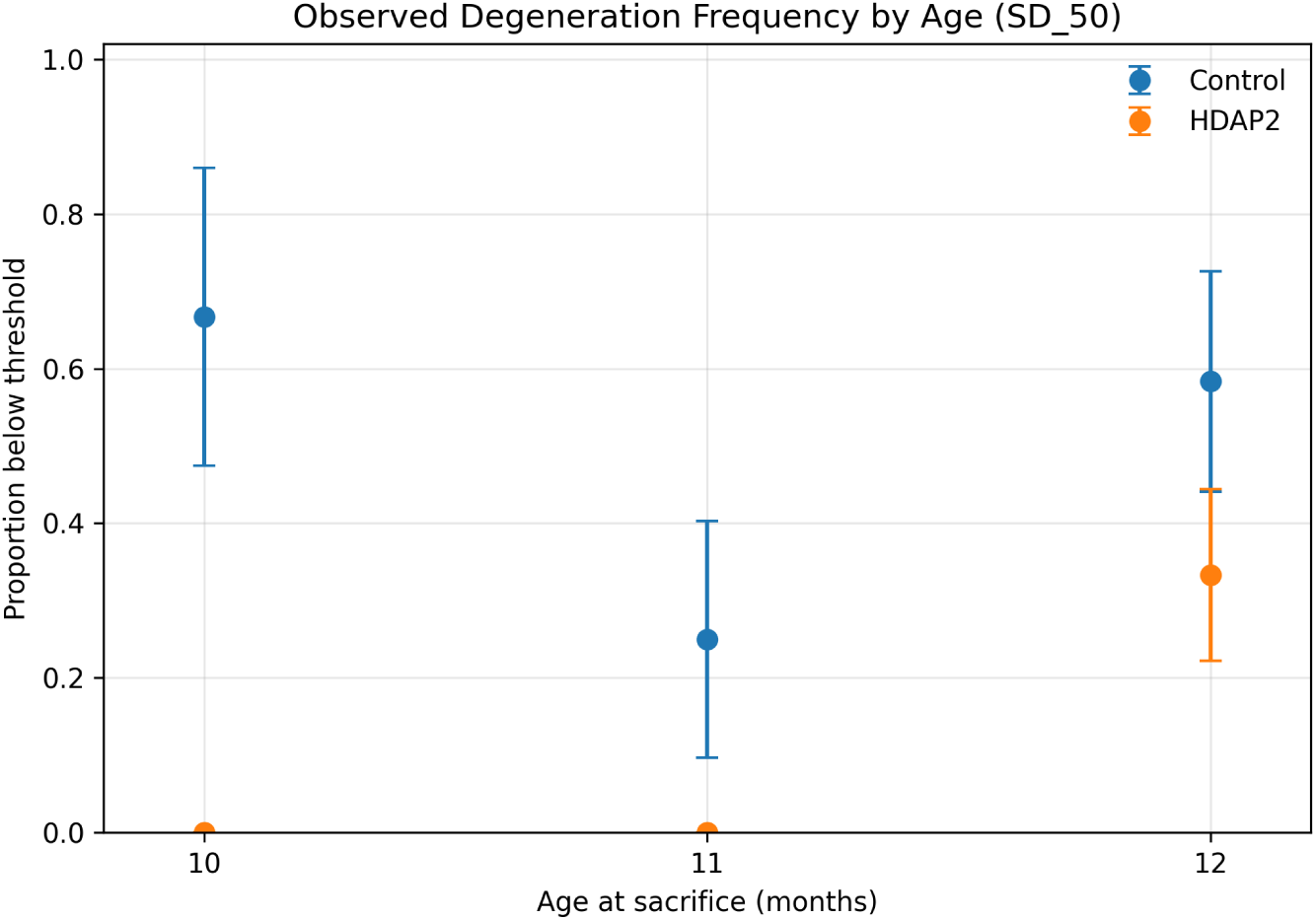
Observed degeneration frequency at the SD_50_ threshold (severe damage).

**Figure 4:**
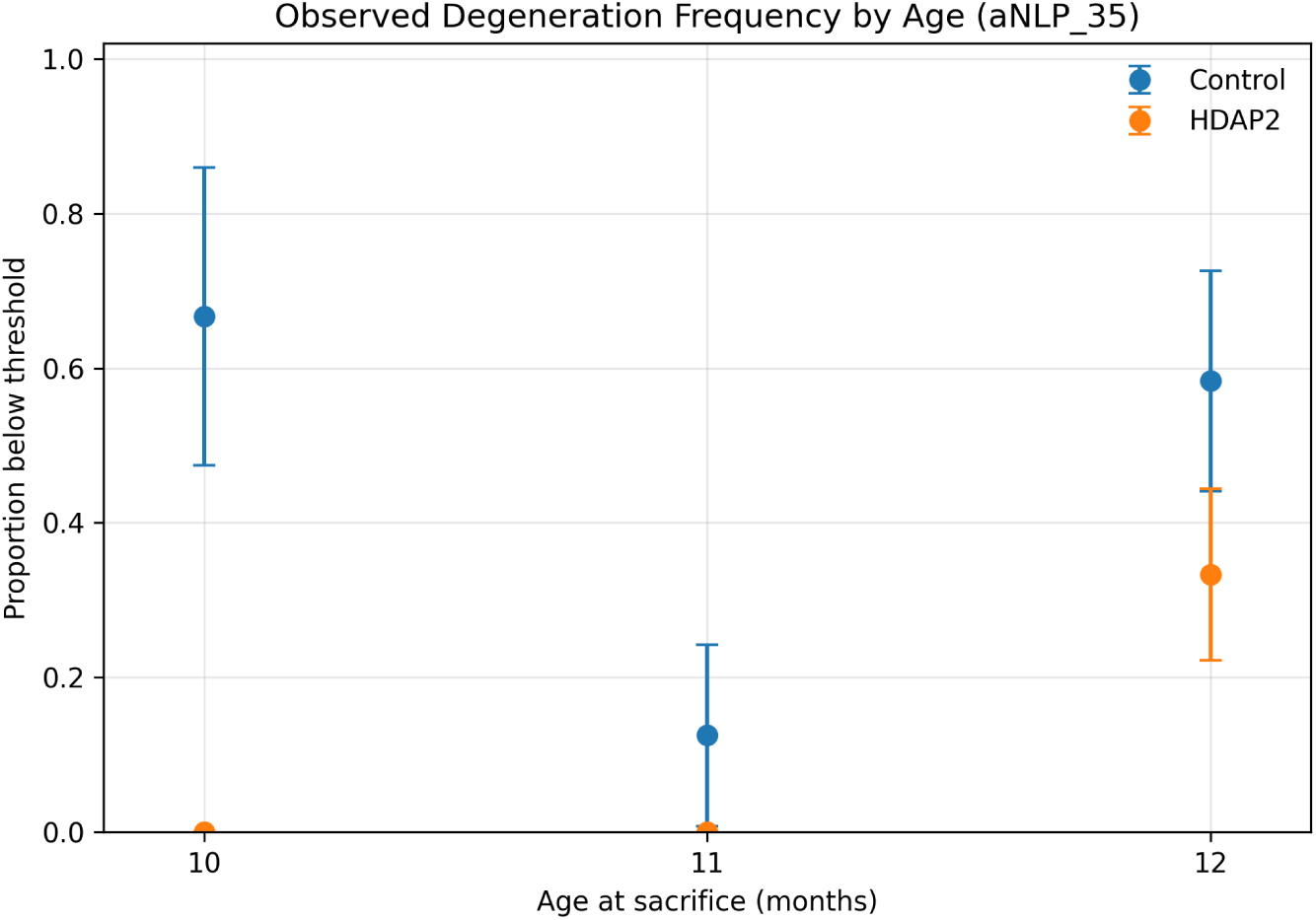
Observed degeneration frequency at the aNLP_35_ threshold (advanced near-complete loss).

**Figure 5:**
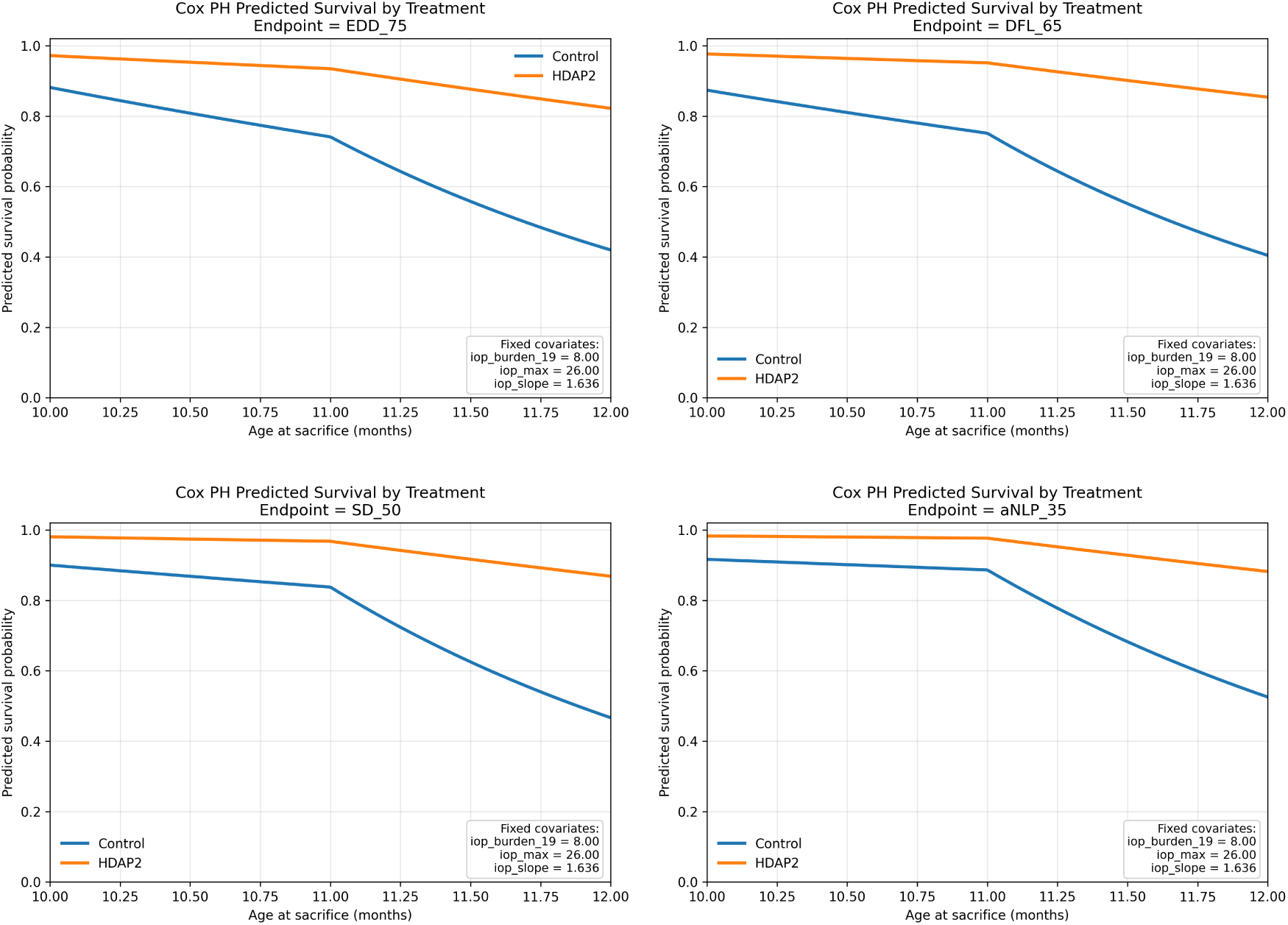
Model-implied survival curves from the proportional hazards model across four RGC loss thresholds (EDD_75_, DFL_65_, SD_50_, and aNLP_35_). For each endpoint, curves are shown for untreated and HDAP2-treated animals under fixed covariate values.

For the same IOP exposure profile, HDAP2 treatment is associated with a consistently higher probability of remaining above each RGC loss threshold at all time points. The separation between treatment and control curves increases over time. This pattern is stable across all endpoints, indicating that the treatment effect is observed across multiple levels of RGC loss severity.

## 5 Time-Dependent Cox Model

The proportional hazards model developed in Section 4 demonstrates that cumulative IOP burden is the dominant predictor of RGC degeneration but represents each animal’s pressure history through a small number of static covariates. This approach provides interpretable estimates of risk but does not distinguish between trajectories that accumulate similar pressure burdens through different temporal patterns. We therefore extend the model to incorporate the evolving IOP trajectory directly, allowing progression risk to depend on both current pressure and accumulated exposure.

We employ a *time-dependent Cox proportional hazards model* in the Andersen–Gill formulation [3], [1], in which each animal contributes multiple observation intervals and the hazard depends on both instantaneous and cumulative exposure.

A joint longitudinal-survival model provides an alternative framework for linking biomarker trajectories to survival outcomes, particularly when measurements are irregular or subject to substantial measurement error [23]. In the present study, IOP was measured monthly under standard conditions, and the inferential target was progression risk conditional on the observed pressure history. The Andersen-Gill formulation therefore provides a simpler and more transparent approach while allowing the observed IOP trajectory to enter the hazard directly.

### 5.1 Interpretation of Time and Events

As in the proportional hazards model, the event is defined by whether a given RGC loss threshold has been reached at the age of sacrifice.

Accordingly, the time-dependent model describes the *intensity of latent threshold crossing* under the observed IOP trajectory rather than the exact timing of the biological event. The framework thus provides a continuous-time representation of risk driven by longitudinal exposure while remaining consistent with the discrete observation structure of the data.

### 5.2 Construction of the Time-Varying Covariates

For each animal *i*, let 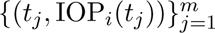 denote the observed IOP measurements. We partition the observation window into intervals [*t_j_, t_j_*_+1_), with the final interval truncated at the sacrifice age, following the standard counting-process representation for time-varying covariates [3], [1]. We define the time-dependent covariates as

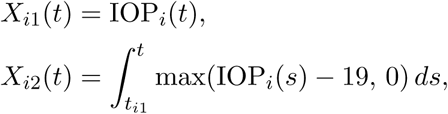

where *X_i_*_1_(*t*) represents the contemporaneous IOP level and *X_i_*_2_(*t*) the cumulative burden above 19 mmHg. Thus, the model incorporates both:

i. an *instantaneous (acute)* effect of IOP, and
ii. a *cumulative (chronic)* effect of sustained elevation.

In practice, these quantities are approximated on each interval [*t_j_, t_j_*_+1_) by

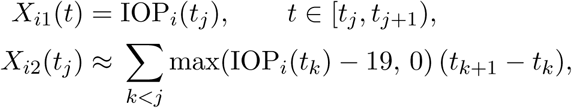

corresponding to a left-Riemann sum approximation.

### 5.3 Model Specification and Estimation

The hazard is modeled as

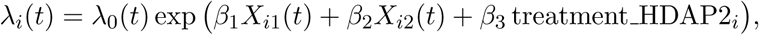

where *λ*_0_(*t*) is a common baseline hazard. The coefficients *β* admit the interpretation:

1. *â*_1_: contemporaneous IOP effect,
2. *â*_2_: cumulative burden effect,
3. *â*_3_: treatment effect (HDAP2).

The model is estimated via the Andersen–Gill partial likelihood [3], [1] using the Python function lifelines CoxTimeVaryingFitter [6], with mild *L*_2_-regularization (penalizer = 0.01) applied to stabilize estimation given the small sample size. Coefficient estimates were not sensitive to penalizer values in the range [0.001, 0.1].

### 5.4 Results

Table 2 summarizes the estimated coefficients across all RGC loss thresholds.

**Table 2:** Time-dependent Cox regression results across RGC loss thresholds. The model includes contemporaneous IOP, cumulative IOP burden above 19 mmHg, and treatment (HDAP2). Hazard ratios (HR) are reported as exp(*β̂*).

| Covariate | $\hat{\beta}$ | SE | *p*-value | HR | 95% CI |
| --- | --- | --- | --- | --- | --- |
| **EDD_75** |  |  |  |  |  |
| iop_current | 0.024 | 0.034 | 0.472 | 1.02 | (0.96, 1.09) |
| cum_burden_19 | 0.111 | 0.027 | 4.63e-05 | 1.12 | (1.06, 1.18) |
| treatment_HDAP2 | -1.291 | 0.422 | 0.00221 | 0.28 | (0.12, 0.63) |
| **DFL_65** |  |  |  |  |  |
| iop_current | 0.026 | 0.034 | 0.453 | 1.03 | (0.96, 1.10) |
| cum_burden_19 | 0.110 | 0.028 | 8.52e-05 | 1.12 | (1.06, 1.18) |
| treatment_HDAP2 | -1.389 | 0.433 | 0.00134 | 0.25 | (0.11, 0.58) |
| **SD_50** |  |  |  |  |  |
| iop_current | 0.038 | 0.036 | 0.288 | 1.04 | (0.97, 1.11) |
| cum_burden_19 | 0.105 | 0.030 | 0.000447 | 1.11 | (1.05, 1.18) |
| treatment_HDAP2 | -1.327 | 0.455 | 0.00352 | 0.27 | (0.11, 0.65) |
| **aNLP_35** |  |  |  |  |  |
| iop_current | 0.053 | 0.036 | 0.138 | 1.05 | (0.98, 1.13) |
| cum_burden_19 | 0.112 | 0.030 | 0.000238 | 1.12 | (1.05, 1.19) |
| treatment_HDAP2 | -1.279 | 0.464 | 0.00588 | 0.28 | (0.11, 0.69) |

The baseline cumulative hazard Λ̂_0_(*t*) is estimated within the Andersen–Gill framework using a Breslow-type estimator applied to the interval-based data [1]. The estimated baseline hazard provides the basis for converting trajectory-dependent relative risk into absolute survival probabilities.

The time-dependent specification reproduces the qualitative conclusions of the static model while representing the IOP trajectory dynamically. The cumulative-burden coefficient remains positive and significant at every threshold (HR *≈* 1.11–1.12), with magnitude and significance closely matching Table 1. In contrast, the contemporaneous IOP coefficient is small and non-significant once cumulative burden is included, indicating that current pressure provides little additional predictive information beyond accumulated exposure.

HDAP2 remained strongly protective across all thresholds, with hazard ratios consistently below unity, indicating reduced progression risk after adjustment for both instantaneous and cumulative IOP exposure.

### 5.5 Model-Implied Survival Comparisons

To visualize the treatment effect, we evaluate the fitted model at representative covariate values (median contemporaneous IOP and cumulative burden) and vary only the treatment indicator.

Across all thresholds, HDAP2 treatment produces a clear rightward shift in the model-implied survival curves, corresponding to delayed attainment of RGC loss endpoints. The treatment effect is consistent with the experimental findings reported in [17], while the model-implied curves extend those observations by representing treatment-associated differences continuously across the range of IOP exposures and sacrifice ages.

### 5.6 Hypothetical IOP Trajectories

A key advantage of the time-dependent formulation is the ability to evaluate survival under prescribed IOP trajectories.

The resulting curves exhibit a clear monotonic ordering, with higher sustained IOP levels associated with systematically accelerated progression.

### 5.7 Trajectory-Based Risk and Relation to the Static Model

The time-dependent Cox model induces a natural mapping from longitudinal IOP trajectories to survival probabilities. Let

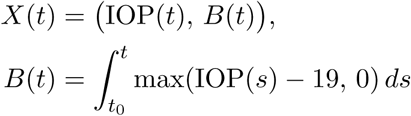

denote the dynamic covariates introduced above. The model implies

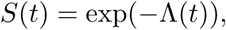

with cumulative hazard

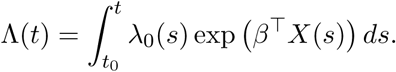

The fitted model thus defines a mapping *F*: IOP(*·*) *−→ S*(*·*) that assigns a survival curve to each admissible IOP trajectory, with cumulative burden *B*(*t*) entering as the dominant functional of the path.

The proportional hazards model of §4 is, in this view, a low-dimensional projection of the same functional: it captures cumulative burden as a scalar summary but discards the temporal information about when exposure accumulated. The two specifications agree on the dominant role of integrated exposure, while the time-dependent formulation additionally shows how this effect evolves over time and permits evaluation under prescribed IOP trajectories (§5.6). Section 6 develops this trajectory-to-survival mapping concretely, reconstructing continuous-time survival curves from the fitted hazard and using them to evaluate hypothetical IOP profiles.

## 6 Model-Implied Survival Curves

The proportional hazards models developed in Sections 4 and 5 quantify how longitudinal IOP exposure influences the hazard of RGC degeneration. To make these results more interpretable, we reconstruct continuous-time, model-implied survival curves by integrating the fitted hazard over prescribed IOP trajectories. These curves represent the probability that an animal has not yet crossed a given RGC loss threshold at each point in its life and can be evaluated under both observed and hypothetical pressure profiles.

Let *λ̂*_0_(*t*) denote the baseline hazard and let *X*(*t*) represent a covariate trajectory (including contemporaneous IOP, cumulative burden, and treatment status). Under the standard relationship between the hazard and survival functions [1], the corresponding estimated survival function is

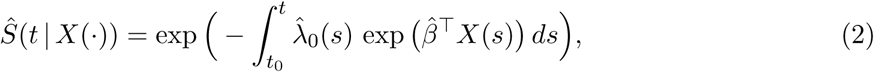

where *t*_0_ = 4 months is the baseline age. This representation defines a trajectory-dependent risk functional that maps longitudinal IOP exposure to model-implied survival.

### 6.1 Reconstruction of the Baseline Hazard

The baseline cumulative hazard Λ̂_0_(*t*) obtained from the Andersen–Gill formulation [3], [1] is defined on a discrete set of event times, reflecting the fact that threshold crossing is only observed at sacrifice ages. Direct use of this estimator produces a step-function hazard and corresponding discontinuities in the implied survival curves.

To obtain a continuous-time approximation, we redistribute the increments of Λ̂_0_(*t*) over the corresponding observation intervals. Specifically, if

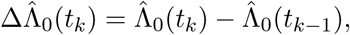

we define a piecewise-constant baseline hazard by

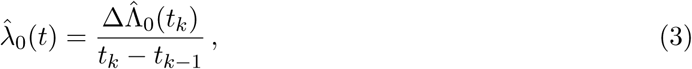

where *t ∈* [*t_k−_*_1_*, t_k_*).

In the present data, this corresponds to distributing the baseline hazard over the intervals [*4, 10*], [*10, 11*], and [*11, 12*] months. This construction preserves the cumulative hazard over each interval while providing a continuous-time representation consistent with the fitted Cox likelihood. We emphasize that this redistribution is a smoothing step used only to construct model-implied survival curves under observed and hypothetical trajectories: the partial-likelihood estimates of *β* reported in Tables 1–2 are unaffected, and the cumulative hazard Λ̂_0_(*t*) is preserved at each observed event time by construction.

### 6.2 Constant IOP Trajectories

Figure 8 shows model-implied survival curves for the primary endpoint DFL_65_ under constant IOP trajectories of 15, 20, and 25 mmHg.

**Figure 6:**
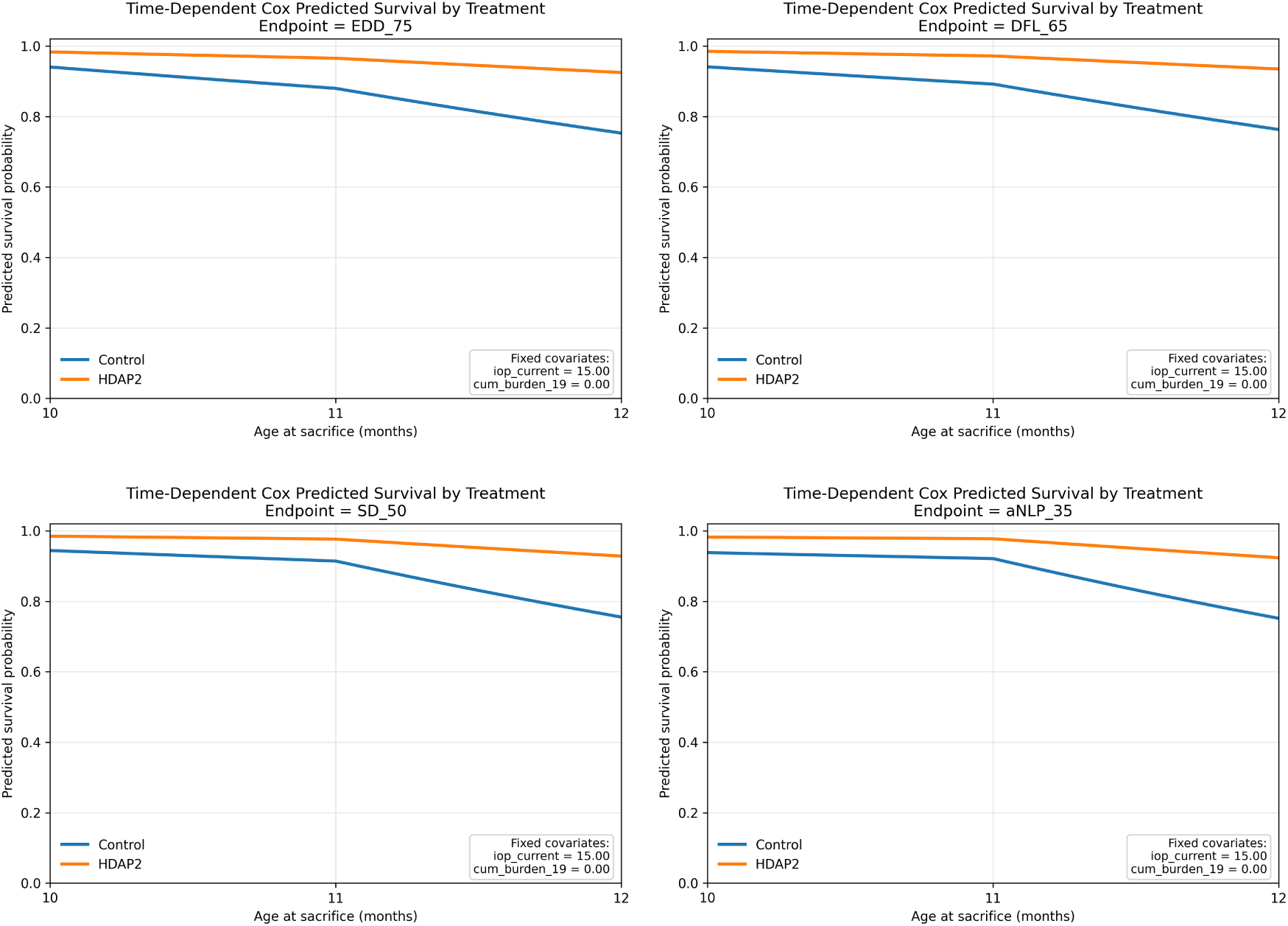
Model-implied survival curves from the time-dependent Cox model comparing untreated and HDAP2-treated animals across RGC loss thresholds. Continuous covariates are fixed at cohort medians. HDAP2 treatment produces a consistent rightward shift in survival curves, indicating delayed progression. This separation is consistent with the hazard ratios reported in Table 2.

**Figure 7:**
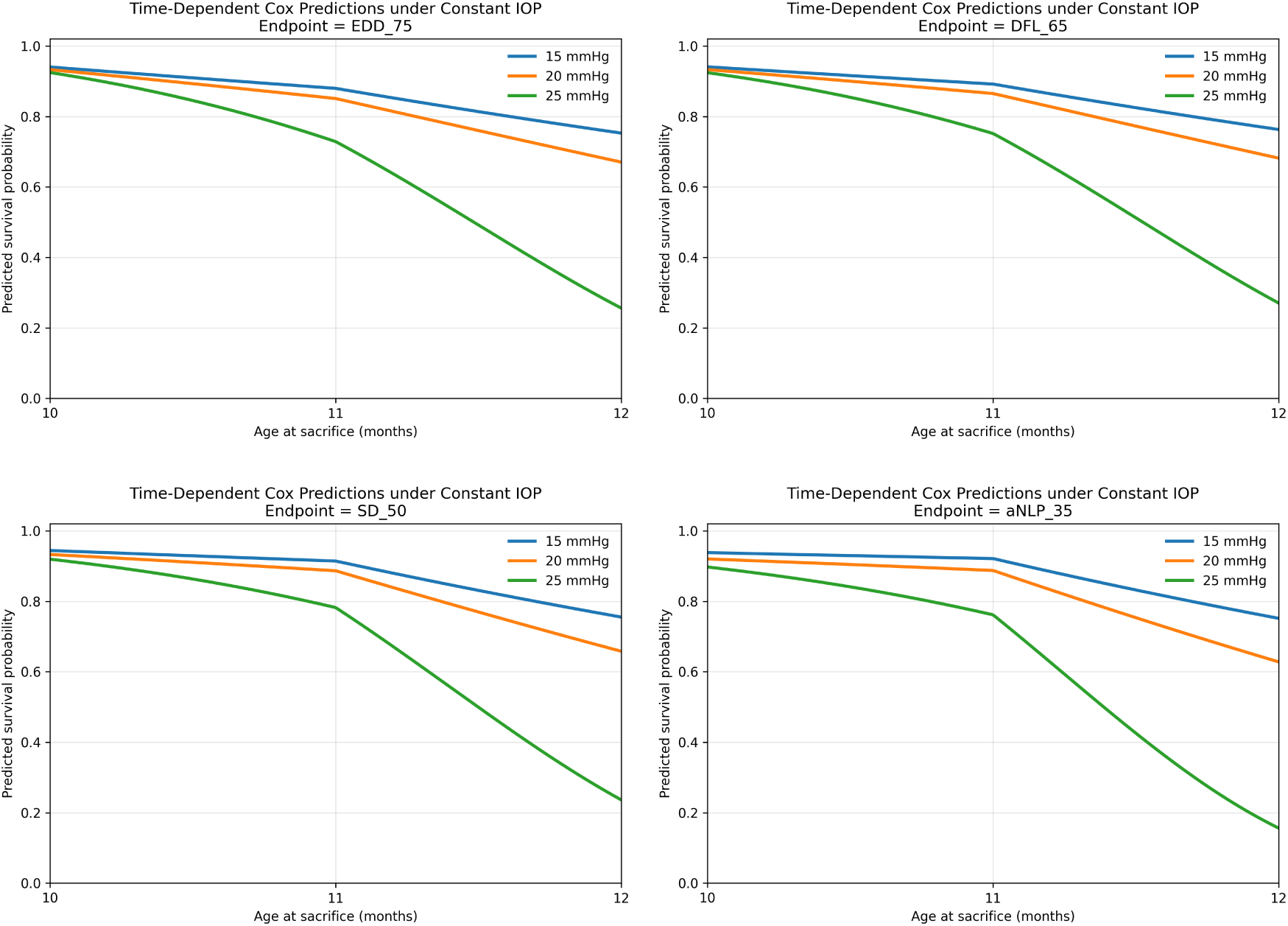
Model-implied survival curves under constant IOP levels (15, 20, 25 mmHg). Higher sustained IOP produces systematically accelerated progression across all thresholds, illustrating the dominant role of cumulative exposure.

**Figure 8:**
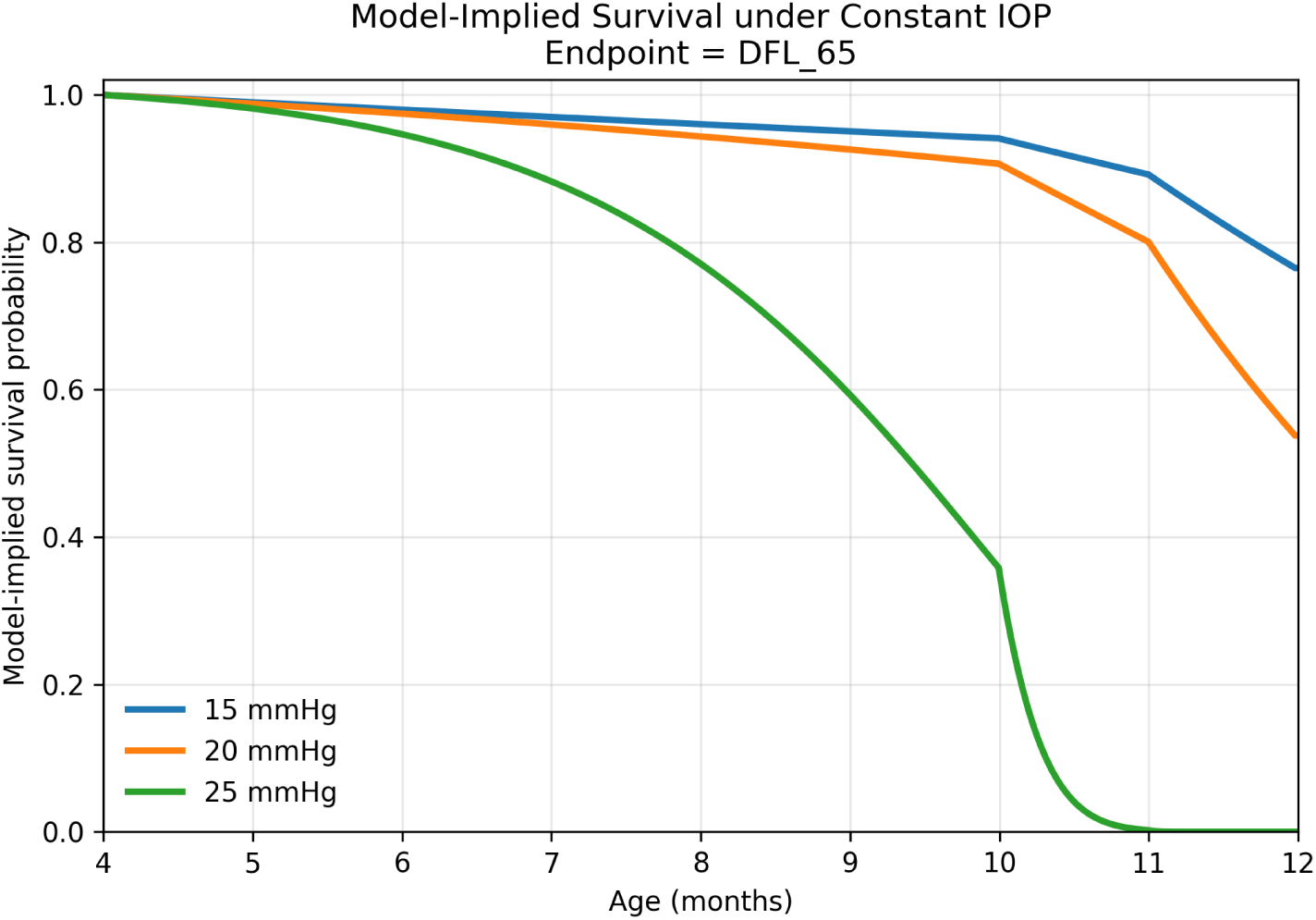
Model-implied survival under constant intraocular pressure (IOP). Survival curves are generated from the fitted time-dependent Cox model using hypothetical constant IOP trajectories (15, 20, and 25 mmHg). The baseline hazard is distributed over observation intervals to obtain a continuous-time approximation. Higher IOP levels lead to progressively earlier and steeper declines in survival, reflecting the cumulative effect of IOP burden.

Figure 8 shows that the model-implied survival curves begin to diverge before the first observed sacrifice age, extending the fitted progression-risk representation into the pre-sacrifice interval.

The timing of progression within this interval is model-implied rather than directly observed. At 25 mmHg, the model-implied survival curve shows a pronounced late decline. Together, these curves illustrate how the fitted model extends the discrete observations into a continuous-time representation of progression risk.

### 6.3 Treatment Effect Under Controlled Exposure

To isolate the effect of treatment from differences in IOP exposure, we evaluate the fitted model under a common prescribed trajectory and compare the resulting survival curves for untreated and HDAP2-treated animals.

Figure 9A shows a rightward shift of the HDAP2 curve relative to the untreated condition under the same IOP exposure profile, indicating delayed progression attributable to treatment. The agreement between panels (A) and (B) provides an internal consistency check: the qualitative pattern of the observed degeneration frequencies at sacrifice ages 10, 11, and 12 months is reproduced by the model-implied latent survival curves derived from the fitted hazard coefficients and baseline hazard estimate. Because both panels are computed from the same underlying data, this agreement does not constitute independent validation; rather it indicates that the reconstruction preserves the qualitative information present in the observed endpoint data.

**Figure 9:**
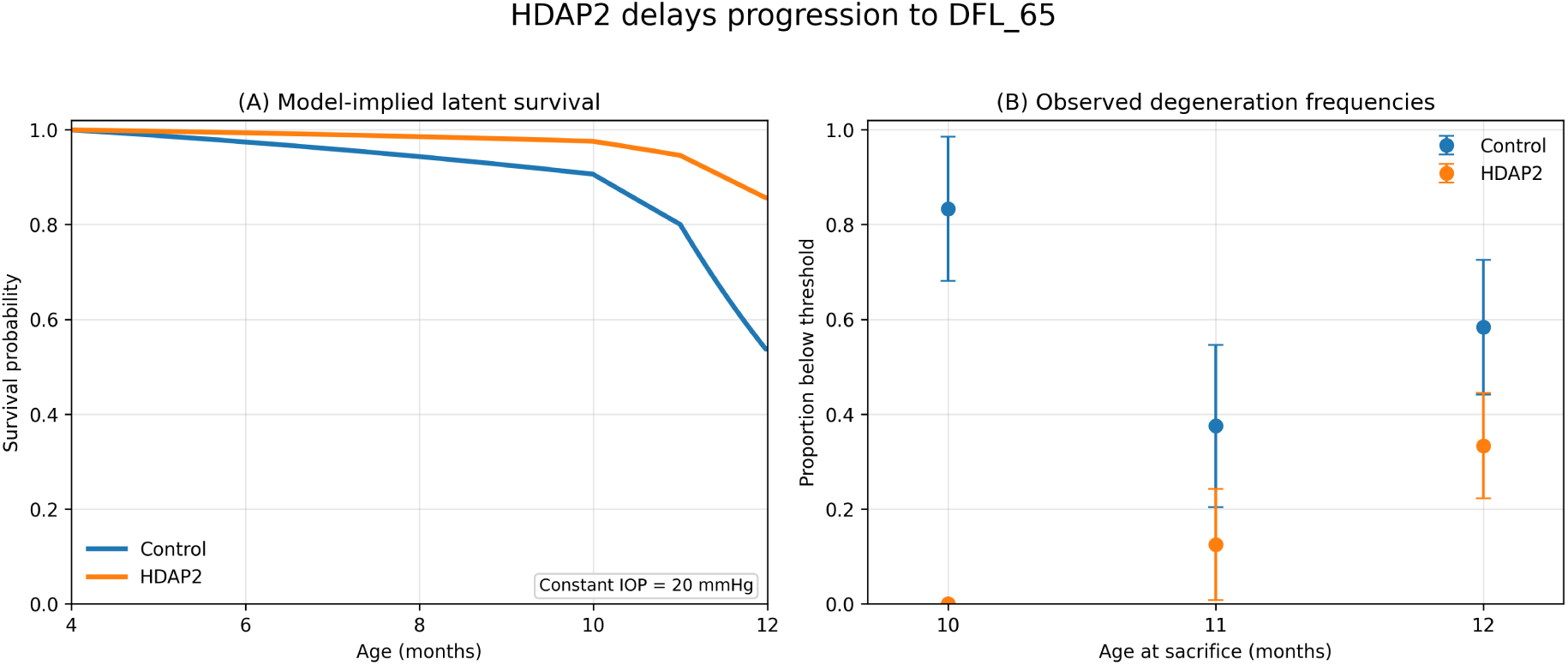
HDAP2 delays progression to the primary endpoint DFL_65_. (A) Model-implied latent survival curves under a common constant IOP trajectory of 20 mmHg, comparing untreated and HDAP2-treated animals. Because the IOP trajectory is identical, the separation between curves isolates the fitted treatment effect. (B) Observed degeneration frequencies at sacrifice ages 10, 11, and 12 months for the same endpoint. The empirical data exhibit the same qualitative pattern, providing an internal consistency check on the model-implied survival dynamics.

## 7 Discussion

This study presents a survival-analysis framework for inferring latent disease progression from longitudinal physiological measurements. Applied to experimental glaucoma, the framework indicates that cumulative intraocular pressure (IOP) burden is the principal predictor of retinal ganglion cell (RGC) degeneration across both proportional hazards and time-dependent Cox models. The agreement between these complementary approaches indicates that cumulative pressure exposure, rather than transient elevations in IOP, is the dominant predictor of disease progression in this experimental model.

These findings extend the original analysis of this experimental cohort [15], which demonstrated that cumulative IOP burden was associated with RGC survival and that HDAP2 shifted the burden threshold for severe degeneration. By incorporating cumulative burden, peak IOP, pressure trajectory, and treatment within a unified hazard framework, the present analysis shows that cumulative burden remains the dominant predictor even after accounting for other features of the longitudinal pressure history. Together, these results support a model in which retinal ganglion cell degeneration reflects the accumulation of chronic physiological stress rather than isolated episodes of elevated pressure.

The time-dependent Cox model further shows that disease progression can be represented as a function of the evolving IOP trajectory rather than a collection of static summary measures. The close agreement between the static and time-dependent analyses indicates that cumulative burden captures the dominant component of the longitudinal pressure history while providing a considerably simpler summary of exposure. More generally, the framework illustrates how longitudinal physiological measurements can be linked to latent biological progression in experimental studies where structural outcomes are observed only at terminal endpoints.

The persistence of the HDAP2 treatment effect after adjustment for longitudinal IOP exposure supports a pressure-independent neuroprotective effect, consistent with a mechanism that enhances RGC resilience to chronic stress. More broadly, the framework provides a quantitative means of separating pressure-dependent and pressure-independent therapeutic effects, offering a useful strategy for evaluating neuroprotective interventions in experimental glaucoma. These findings also contribute to the broader discussion regarding which aspects of longitudinal IOP history best predict glaucoma progression. Within the controlled setting of the DBA/2J model, cumulative IOP burden consistently emerged as the dominant predictor, supporting the view that sustained exposure provides more prognostic information than isolated pressure measurements. Whether this relationship extends to heterogeneous clinical populations remains an important question for future investigation.

This study has several limitations. The experimental cohort is modest in size, resulting in greater uncertainty for some degeneration thresholds, particularly SD50. Sex was not included as a covariate and should be considered in future studies with larger cohorts. Confirmation in independent experimental datasets will be important for establishing the generality of the present findings.

Although developed for experimental glaucoma, the framework is applicable to a broad class of biological problems in which longitudinal physiological measurements precede irreversible biological outcomes. By linking exposure histories to latent survival processes, it provides a principled approach for reconstructing disease progression, evaluating hypothetical interventions, and quantifying therapeutic effects. More broadly, these results suggest that longitudinal physiological measurements often contain substantially more information about disease progression than is apparent from terminal endpoint analyses alone.

## Declaration of generative AI and AI-assisted technologies in the manuscript preparation process

During the preparation of this work, the authors used ChatGPT (OpenAI) and Claude (Anthropic) for language editing and stylistic refinement of portions of the manuscript. All substantive ideas, models, derivations, and conclusions are solely those of the authors. After using these tools, the authors reviewed and edited the content as needed and take full responsibility for the content of the published article.

## References

[1] Andersen, P. K., Borgan, O., Gill, R. D., and Keiding, N.: Statistical Models Based on Counting Processes, Springer-Verlag (1993).

[2] Anderson, M. G., Smith, R. S., Hawes, N. L., Zabaleta, A., Chang, B., Wiggs J. L., and John, S. W. M.: Mutations in genes encoding melanosomal proteins cause pigmentary glaucoma in DBA/2J mice, Nat Genet, 30,: 81–85 (2002).

3. Andersen, P. K. and Gill, R. D.: Cox’s Regression Model for Counting Processes: A Large Sample Study, Annals of Statistics, 10, 1100–1120 (1982).

[4] Asrani, S. G., McGlumphy, E. J., Al-Aswad, L. A., Chaya, C. J., Lin, S., Musch, D. C., Pitha, I., Robin, A. L., Wirostko, B., and Johnson, T. V.: The relationship between intraocular pressure and glaucoma: an evolving concept, Prog. Retin. Eye Res., 103, 101303 (2024).

[5] Calkins, D. J.: Critical pathogenic events underlying progression of neurodegeneration in glaucoma, Prog. Retin. Eye Res., 31, 702–719 (2012).

[6] Davidson-Pilon, C.: lifelines: survival analysis in Python, Journal of Open Source Software, 4(40), 1317, 10.21105/joss.01317 (2019).

[7] De Moraes, C. G., Juthani, V. J., Liebmann, J. M., Teng, C. C., Tello, C., Susanna, R. Jr., and Ritch, R.: Risk factors for visual field progression in treated glaucoma, Arch. Ophthalmol., 129(5), 562–568 (2011).

[8] Harwerth, R. S., Carter-Dawson, L., Shen, F., Smith, E. L., and Crawford, M. L.: Ganglion cell losses underlying visual field defects from experimental glaucoma, Invest. Ophthalmol. Vis. Sci., 40, 2242–2250 (1999).

[9] Hedberg-Buenz, A., Christopher, M. A., Lewis, C. J., Fernandes, K. A., Dutca, L. M., Wang, K., Scheetz, T. E., Abramoff, M. D., Libby, R. T., Garvin, M. K., and Anderson, M. G.: Quantitative measurement of retinal ganglion cell populations via histology-based random forest classification, Exp Eye Res 146, 370–385 (2016).

[10] Heijl, A., Leske, M. C., Bengtsson, B., et al.: Reduction of intraocular pressure and glaucoma progression: results from the Early Manifest Glaucoma Trial, Arch. Ophthalmol., 120, 1268– 1279 (2002).

[11] Holcombe, D. J., Lengefeld, N., Gole, G. A., and Barnett, N. L.: Selective inner retinal dysfunction precedes ganglion cell loss in a mouse glaucoma model, Br. J. Ophthalmol., 92(5), 683–688 (2008).

[12] Hou, K., Bradley, C., Herbert, P., Johnson, C., Wall, M., Ramulu, P. Y., Unberath, M., and Yohannan, J.: Predicting visual field worsening with longitudinal OCT data using a gated transformer network, Ophthalmology, 130(8), 854–862 (2023).

[13] Ishida, K., Yamamoto, T., and Kitazawa, Y.: Clinical factors associated with progression of normal-tension glaucoma, J. Glaucoma, 7(6), 372–377 (1998).

[14] Jakobs, T. C., Libby, R. T., Ben, Y., John, S., and Masland, R. H.: Retinal ganglion cell degeneration is topological but not cell type specific in DBA/2J mice, The Journal of Cell Biology, 171(2), 313–325. 10.1083/jcb.200506099 PMID - 16247030 (2005).

[15] John, S. W., Smith, R. S., Savinova, O. V., Hawes, N. L., Chang, B., Turnbull, D., Davisson, M., Roderick, T. H., and Heckenlively, J. R.: Essential iris atrophy, pigment dispersion, and glaucoma in DBA/2J mice, Invest Ophthalmol Vis Sci, 39, 951–962 (1998).

[16] Libby, R. T., Anderson, M. G., Pang, I.-H., et al.: Inherited glaucoma in DBA/2J mice: pertinent disease features for studying the neurodegeneration, Vis. Neurosci., 22, 637–648 (2005).

17. MacNeil, M. A., Mentor, W., and Birk, A.: Mitochondrial-targeted HDAP2 preserves retinal ganglion cells and increases pressure tolerance in DBA/2J mice, Invest. Ophthalmol. Vis. Sci., 66(15), 10.1167/iovs.66.15.66 (2025).

[18] Musch, D. C., Gillespie, B. W., Lichter, P. R., Niziol, L. M., and Janz, N. K.: Visual field progression in the Collaborative Initial Glaucoma Treatment Study: the impact of treatment and other baseline factors, Ophthalmology, 116(2), 200–207 (2009).

[19] Nagaraju, M., Saleh, M., and Porciatti, V.: IOP-dependent retinal ganglion cell dysfunction in glaucomatous DBA/2J mice, Invest. Ophthalmol. Vis. Sci., 48(10), 4573–4579 (2007).

[20] Nouri-Mahdavi, K., et al.: Predictive factors for glaucomatous visual field progression in the Advanced Glaucoma Intervention Study, Ophthalmology, 111, 1627–1635 (2004).

[21] Porciatti, V., Saleh, M., and Nagaraju, M.: The pattern electroretinogram as a tool to monitor progressive retinal ganglion cell dysfunction in the DBA/2J mouse model of glaucoma, Invest. Ophthalmol. Vis. Sci., 48(2), 745–751 (2007).

[22] Quigley, H. A., Addicks, E. M., and Green, W. R.: Optic nerve damage in human glaucoma. III. Quantitative correlation of nerve fiber loss and visual field defect in glaucoma, ischemic neuropathy, papilledema, and toxic neuropathy, Arch. Ophthalmol., 100, 135–146 (1982).

23. Rizopoulos, D.: Joint Models for Longitudinal and Time-to-Event Data: With Applications in R, CRC Press (Taylor & Francis Group), 10.1201/b12208 (2012).

[24] Shin, Y. I., Jeong, Y., Huh, M. G., Kim, Y. K., Park, K. H., and Jeoung, J. W.: Longitudinal evaluation of advanced glaucoma: ten year follow-up cohort study, Scientific Reports, 14(1), 476 (2024).

[25] Tao, S., Ravindranath, R., and Wang, S. Y.: Predicting glaucoma progression to surgery with artificial intelligence survival models, Ophthalmology Science, 3(4), 100336 (2023).

[26] Turner, A. J., Vander Wall, R., Gupta, V., Klistorner, A., and Graham, S. L.: DBA/2J mouse model for experimental glaucoma: pitfalls and problems, Clin. Exp. Ophthalmol., 45(9), 911– 922 (2017).

